# Cell viability assessment associated with a contact of gas bubbles produced by femtosecond laser breakdown in cell culture media

**DOI:** 10.1101/2022.01.16.473385

**Authors:** Ryohei Yasukuni, Akari Koyanagi, Yukihiro Tanaka, Kazunori Okano, Yoichiroh Hosokawa

## Abstract

High intensity near infrared femtosecond laser is a promising tool for three-dimensional processing of biological materials. During the processing of cells and tissues, long lasting gas bubbles randomly appeared around the laser focal point, however physicochemical and mechanical effects of the gas bubbles has not been emphasized. This paper presents characteristic behaviors of the gas gabbles and their contact effects on cell viability. High-speed imaging of the gas bubble formation with various additives in physiological medium confirms that the gas bubble consists of dissolved air, and amphipathic proteins stabilize the bubble surface. This surface protective layer reduces interactions of gas bubbles and cell membranes. Consequently, the gas bubble contact does not cause critical effects on cell viability. On the other hands, burst of gas bubbles stimulated by an impact of femtosecond laser induced cavitation can lead to liquid jet flow that might cause serious mechanical damages on cells. These results provide insights for the parameter of biological tissue processing with intense fs laser pulses.

## 1. Introduction

High intensity near infrared (NIR) femtosecond laser (fs laser) is a promising tool for three-dimensional processing of transparent biological materials without causing sever thermal damages [1–3]. When the intense NIR fs laser pulse is focused into a biological medium through an objective lens, non-linear processes represented by multiphoton absorption efficiently lead to breakdown and plasma formation [4]. The plasma relaxation within a timescale of thermal diffusion causes rapid elevation of thermodynamic stress at the laser focal point, which has a large contribution to non-thermal characteristics of the fs laser processing [5, 6]. The plasma relaxation in aqueous medium is associated with the occurrence of cavitation bubble formation [4]. Size of the cavitation bubble ranges from hundreds of nanometers to tens of microns, and the bubble repeats expansion and contraction with a frequency up to around 1 MHz with a common laser pulse energy for biomaterial processing [7].

Previous works in our group shows various applications based on the fs laser breakdown in water. We especially exploited the propagation of impulsive hydrodynamic force to the periphery associated with the rapid expansion of the cavitation bubble. Since this hydrodynamic force possesses good spatiotemporal reproducibility and controllability, it enables to evaluate mechanical properties in cells and tissues, for example extrusion forces and surface tension modifications in zebrafish embryonic epithelia in developmental biology [8, 9]. Single cell separation in high-speed fluidics using the fs laser induced hydrodynamic force has also demonstrated toward an ultrafast cell sorting system [10]. Following contraction of the cavitation bubble, micro-sized gas bubbles often remain around the laser focal point. A contact of this fs laser breakdown-induced gas bubbles to cells and tissues was also confirmed under an optical microscope in above applications [8–10].

In an application to the laser vision correction using amplified fs laser pulses, named femto-LASIK, a long-lasting characteristic of the micro gas bubbles induced by fs laser breakdown is used for a flap creation by connecting gas bubbles in a plane within a cornea [11, 12]. The fs laser assisted technique allows more accurate and safer flap creations for various cornea shapes compared to the traditional microkeratome. However, unlike the cavitation bubbles, gas bubbles are randomly appeared thus difficult to control its locations, sizes and lifetimes precisely. The gas bubbles confined to the corneal stroma during a flap creation can disperse spontaneously or be removed by manual surgical techniques. By contrast, an opaque gas bubble layer accidentally forms inside the anterior eye chamber has a potential risk of complications [11, 12].

More commonly in biomedical field, micrometer sized gas bubbles have been used as contrast agents for ultrasound imaging [13, 14]. The bubbles of low solubility and non-toxic gas are stabilized by proteins, lipids or polymers, and they are injected into a blood vessel [13]. Ultrasound pressure waves induce volumetric bubble oscillations that produces large scattering acoustic signal, and it enhances the ultrasound image contrast. The stabilized gas bubble itself has basically no interaction with vascular endothelial cells thanks to their surface protective molecular shell. On the other hand, bubble collapse beside cells, for example by high-pressure ultrasound excitation, leads to disruption and temporal permeability increase of a cell membrane. This effect is known as sonoporation and applied for gene introduction and drug delivery [14–16].

The effects of gas bubbles larger than a few tens of micron have also been investigated from viewpoints of decompression sickness and gas embolism within a blood vessel. In these cases, gas bubbles are originated from a rapid decreasing in the pressure during diving and an injection during surgical procedures. Walsh et al. demonstrated that cell viability decreased when cells were exposed to oxygen bubbles formed by decompression in a 3D engineered tissue phantom [17]. Sobolewski et al. has reported the contact of an air bubble on endothelial cells mechanically induced intracellular calcium transients associated with cell injury and death [18, 19]. In these reports, the large gas bubbles have more of negative effects on a cell viability.

As noted above, the interaction of gas bubbles with cells and tissues depends on various factors such as the gas bubble sizes, surface conditions, environments and external stimuli. Therefore, for further use of fs laser in biomedical applications, specific physicochemical behaviors of gas bubbles induced by fs laser breakdown should be clarified.

In this article, we investigated physicochemical effects of a gas bubble contact on cells in a framework of fs laser applications in biomedical field. We first discussed contents and formation mechanisms of the gas bubble by fs laser break down in cell culture mediums from the results of high-speed imaging in different environments. Then, effects of the gas bubble contact on cultured cells were evaluated in terms of cell viability. Moreover, the effects of gas bubbles were extended from single to multiple fs laser pulses to take account of cavitation-gas bubble interactions.

## 2. Materials and methods

### 2.1 Cell culture

Mouse myoblast cell line (C2C12, RCB0987) was purchased from the RIKEN BRC, Japan. The cells were cultured in Dulbecco’s Modified Eagle Medium (DMEM, Nacalai tesque) supplemented with 10 % of fetal bovine serum (FBS, Invitrogen) and antibiotic agents (100 units/mL of penicillin and 100 μg/mL of streptomycin, Nacalai tesque) using standard cultivation conditions (37°C, 5% CO_2_). For microscope experiments, the cells were seeded on a φ35 cell culture dish with a φ12 glass base (IWAKI). The cell line within 20 passages was used to keep original phenotype.

### 2.2 Gas bubble formation and imaging

Gas bubble formation was performed with a femtosecond Ti:sapphire laser amplifier (Spectra-Physics, Solstice-Ref-MT5W, 800 nm, 150 fs) coupled with an inverted microscope (Olympus, IX71) as shown in Fig. 1. A single femtosecond laser pulse was introduced into the microscope and focused through a 20x objective lens (Olympus, NA: 0.46) in aqueous medium at 10 μm above the image plane. The laser pulse was extracted using a mechanical shutter with a gate time of 1/32 s from 32 Hz pulse trains. The laser pulse energy was tuned with a combination of half waveplate and polarizer, and neutral density filter, and it was measured with a laser power meter (Ophir, Nova Display-Rohr) after the objective lens. The cell culture dish was placed on a motorized microscope stage (Sigma Koki, E-65GR) equipped on the microscope. Gas bubbles with lifetimes less than 50 ms, and over 50 ms were observed with a high-speed camera (Phantom V, Nobby Tech. Ltd.) and a CMOS camera (ORCA-Frash3.0, Hamamatsu) respectively.

**Fig. 1.**
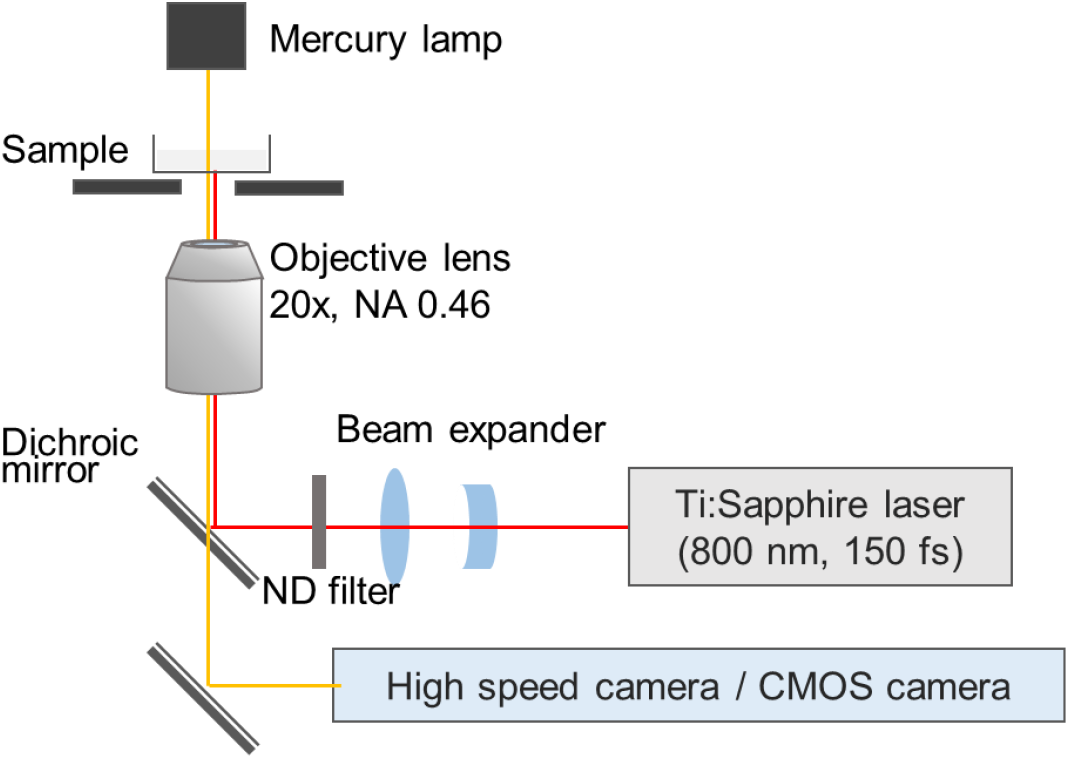
Schematic illustrations of the femtosecond laser induced gas bubble generation and observation system coupled with the inverted optical microscope.

Degassed distilled water and PBS were prepared by following protocols. Both solutions were heated in a microwave oven at 750 W for 1 min, and sonicated at 50°C for 5 min. These steps were repeated three times. Then, the solutions were kept in a vacuum chamber.

Lifetimes of the fs laser breakdown generated gas bubble were compared in the DMEM, in 10% Dulbecco’s Phosphate Buffered Saline (PBS, Nacalai tesque), in the PBS with 5 mg/mL of sterile-filtered bovine serum albumin (BSA, Sigma Aldrich) solution or with 5 mg/mL glucose solution (Sigma Aldrich).

### 2.3 Cell viability assessment after gas bubble contact

Cell viability was assessed by staining dead cells using trypan blue solution (Nacalai tesque). The cell culture medium C2C12 cells were cultured on a glass grid plate (Matsunami) placed in the cell culture dish to identify which cells had a contact with gas bubbles.

## 3. Results and discussion

### 3.1 Characteristic behaviors of the gas bubble

Representative high-speed transmission images of the gas bubble formed in DMEM is shown in Fig. 2a. Along with an appearance of a cavitation bubble at a laser focal point, formation of a gas bubble was confirmed in the cavitation bubble. After disappearing of the cavitation bubble, the micro gas bubble remained near laser focal point. Hereafter this remained micro gas bubble is mentioned just as the gas bubble in distinction from the cavitation bubble.

**Fig. 2.**
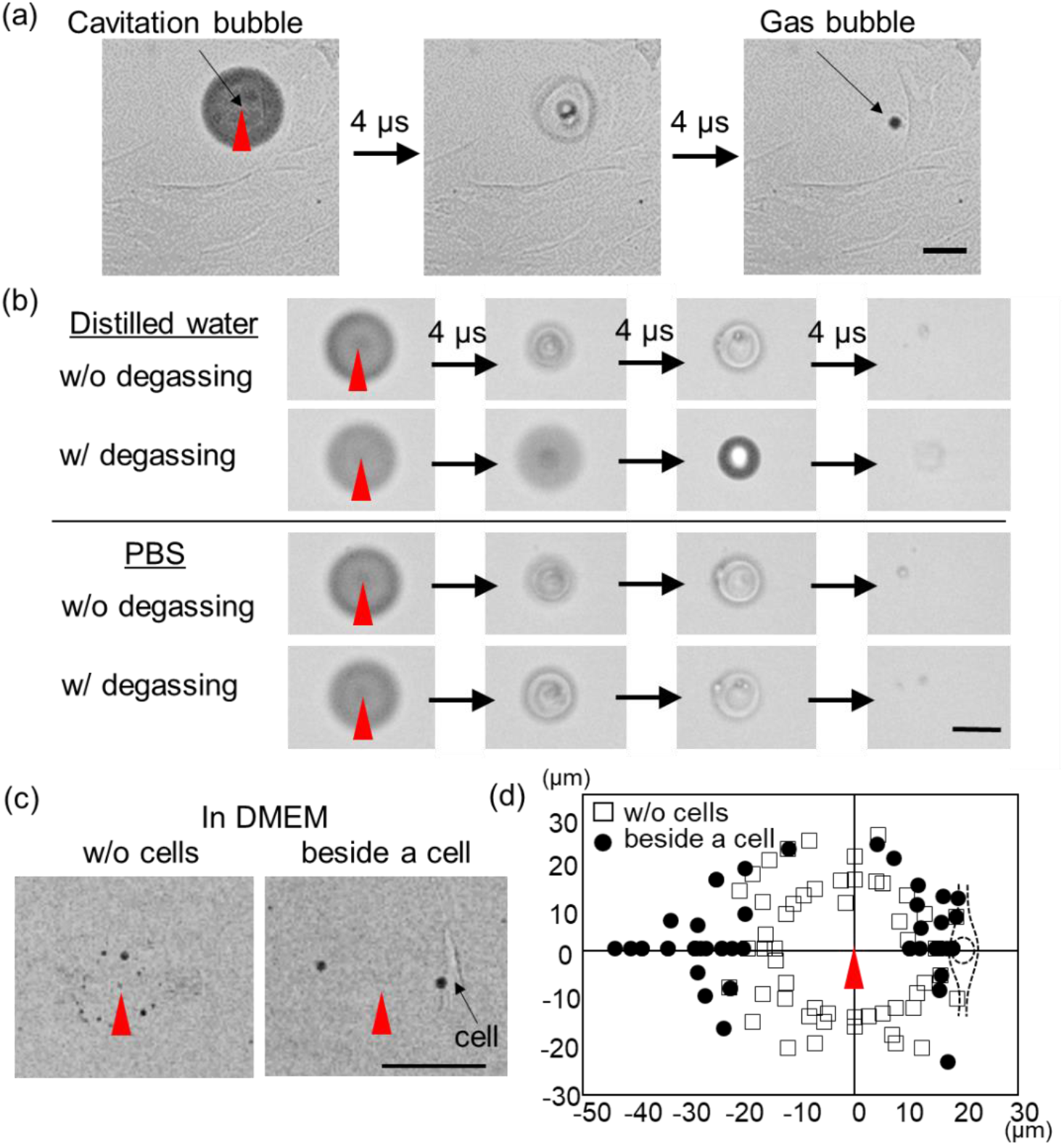
fs laser breakdown induced gas bubble generation observed by the high-speed camera. Single fs laser pulse with 300 nJ was used. (a) A series of high-speed transmission images of the gas bubble forming in DMEM at 4 μs interval. (b) Effects of the degassing on the gas bubble forming in distilled water and in PBS. (c) The high-speed transmission images of the gas bubble formation in DMEM without cells and beside cells. (d) A summary of the bubble remaining positions for without a cell and beside a cell in DMEM. The cell position relative to the laser focal point is also illustrated. The scale bars are 50 μm for all images. The red triangles indicate the laser focal point.

The gas bubble contents were investigated by comparing an effect of degassing in distilled water and in PBS. As shown in Fig. 2b, the gas bubble appeared in distilled water without degassing while it did not appear after degassing. Therefore, the main gas content could be dissolved air in distilled water, which results from decompression through the contraction of the cavitation bubble. In PBS, the gas bubble always appeared regardless of the degassing treatment, which indicated photochemical decompositions of the dissolved molecules also existed in the gas.

Next, the location where the gas bubble remains are examined. The gas bubble remaining positions observed by the high-speed imaging are compared when the laser is focused in DMEM without cells and beside cells as shown in Fig. 2c. Multiple small bubbles tended to remain in all directions around the laser focal point in solution without cell. On the other hand, relatively large two bubbles were more likely remained on the line connecting the laser focal point and the cell when the laser was focused close to cells. This tendency of the gas bubble remaining positions is summarized in Fig. 2d for 15 independent laser shots in both conditions. The observed aspherical gas bubble remaining position could be explained by the cavitation bubble behavior near boundary. Aspherical collapse of the cavitation bubble develops liquid jet flow due to presences of a solid glass substrate and a soft cell membrane, that would align the gas bubbles as seen in Fig. 2d.

A lifetime of the gas bubbles at room temperature are investigated in four mediums; DMEM, PBS, glucose added PBS (PBS-Glu) and BSA added PBS (PBS-BSA). Concentrations of the glucose and BSA were set based on a protein concentration of 10% FBS in DMEM. A single fs laser pulse with 300 nJ was focused into the solutions, and the lifetime (τ) of the gas bubbles was measured with either high speed camera or CMOS camera depending on the lifetime. One shot of the laser pulse gives several gas bubbles in the mediums, and then the maximum τ among the generated bubbles was averaged (τ_ave_) for 10 independent laser irradiations. τ_ave_ acquired in different mediums are shown in Fig. 3. τ_ave_ has similar value in PBS and PBS-Glu, and they are 167 ms and 133 ms, respectively. τ_ave_ in DMEM is 2.04 s and in PBS-BSA is 26.2 s, which are more than 10 times longer than in PBS and PBS-Glu. Bubble lifetimes are generally related to their surface conditions. Surface protective layer stabilize the bubble surface, resulting in lifetime increase. Indeed, initially developed contrast agents for ultrasound imaging were stabilized using human albumin [13]. Therefore, it is considered amphipathic proteins in the solutions work as a protective layer of the bubble surface. Different bubble lifetimes in DMEM and in PBS-BSA would arise from different affinities of FBS in DMEM and BSA in PBS for the bubble surface.

**Fig. 3.**
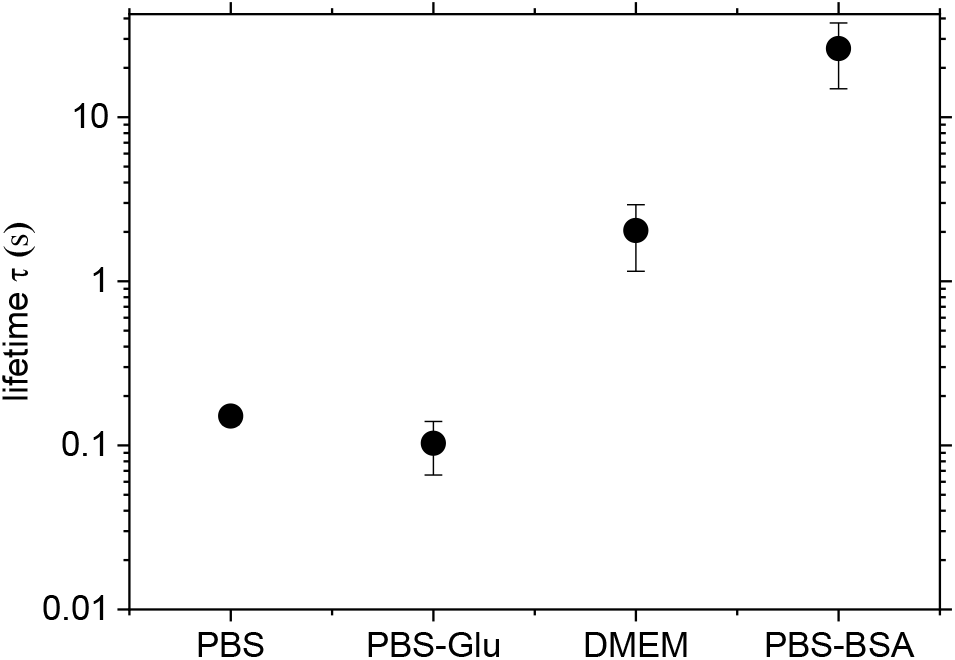
Lifetimes of the gas bubble in aqueous solutions with different solutes. The error bar shows standard deviation of the lifetime obtained from 10 bubbles each.

### 3.2 Effect of the gas bubble contact on cell viability

The results above show that the gas bubbles remain more frequently near the cells and they may interact with lipids and proteins of cell membrane. To understand physiological effects of a gas bubble contact, cell viability after fs laser irradiation near cells was evaluated. The fs laser pulse energy was set at 200, 250 and 300 nJ/pulse and focused 10 μm distance from a target cell. As shown in Fig. 4a, cells were cultured in DMEM on a glass plate with a grid to identify which cells had a contact with the gas bubbles. Cell viability was evaluated from the trypan blue staining by exchanging DMEM for the trypan blue solution. The number of stained cells by trypan blue was summarized in Fig. 4b. The cases of bubble contact and non-contact were visually judged from transmission images with the CMOS camera just for the purpose of reference. At least 9 of 20 cells got a contact with the gas bubble when 200 nJ/pulse was used, and no cell were stained with trypan blue. A number of contact case increased when the pulse energy was 250 nJ, however only one cell was stained and it was a non-contacted case. Four cells in 13 bubble contacted cells were stained when the pulse energy was 300 nJ. Overall, we conclude that the bubble contact itself did not cause a critical effect on cell viability because most of bubble-contacted cells were not stained. This result is supported by the observation in the section 3.1 that bubble surface would be covered by amphipathic molecules immediately after bubble generation, which inhibits chemical interactions with a cell membrane. It is noted that effects of the gas bubble contact and the mechanical force driven by the cavitation bubble expansion should be separately considered. The increased staining rate at the pulse energy of 300 nJ/pulse is regarded as intense mechanical stimuli reduce the cell membrane integrity. We indeed exploited such mechanical force for molecular introduction to a tobacco bright yellow 2 cell through formation of a cleavage on its cell membrane [20].

**Fig. 4.**
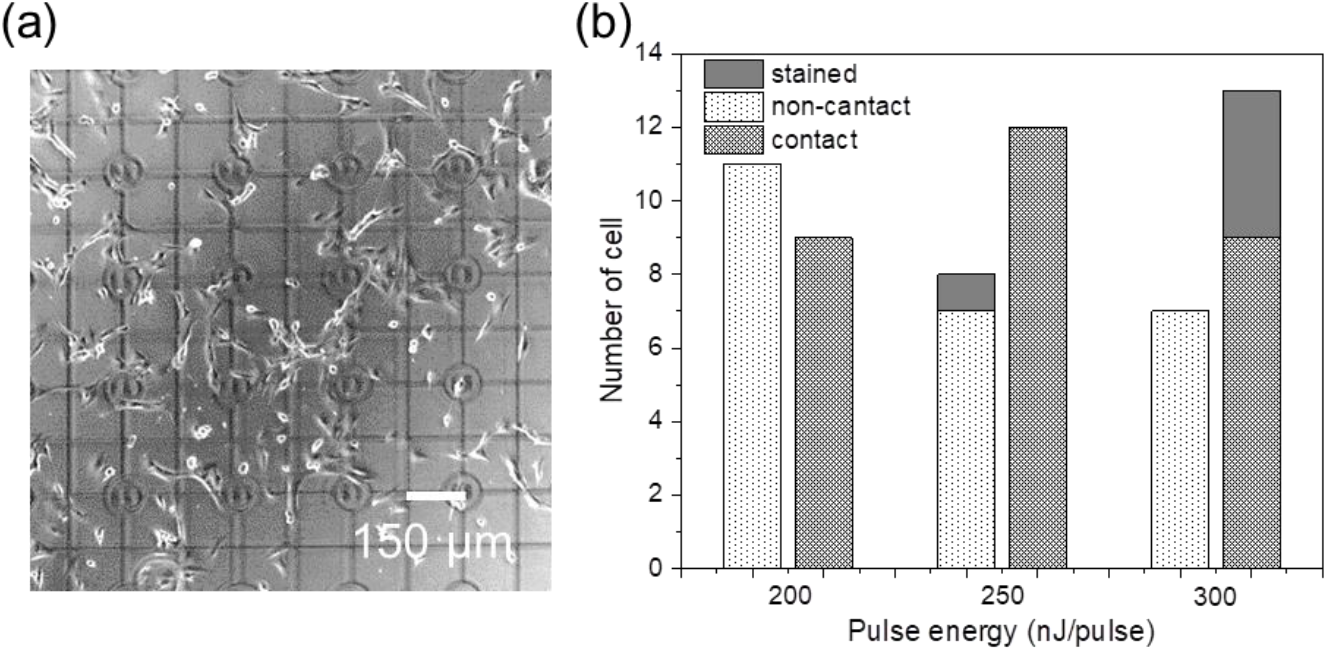
(a) Transmission images of C2C12 cultured on a cover glass with a 150 μm grid. (b) A histogram of the number of trypan blue stained cells after the fs laser irradiation with different pulse energies.

### 3.3 Interaction between cavitation and gas bubbles

Interactions of a cavitation bubble with another cavitation bubble or a gas bubble have been reported previously [21–23]. Such bubble interactions led to a jet flow from one bubble to the other, and this flux enable to a pore formation on a cell membrane [24]. As described above, we found that the lifetime of the gas bubbles is more than 2 sec in DMEM. Hence, when fs laser pulses are focused into a medium over 0.5 Hz, the remained gas bubble can interact with the cavitation bubble induced by the following fs laser pulses. In this context, the gas bubble behavior under sequential fs laser pulses were investigated.

As displayed in Fig. 5, the gas bubbles remained after contraction of the cavitation bubble induced by the first fs laser pulse with 300 nJ focused into the cell culture medium. These gas bubbles interacted with another cavitation bubble induced by a successive fs laser pulse, and larger bubbles appeared around the cavitation bubble. Once the gas bubble reached a certain size, the gas bubble had burst by the impact of the cavitation bubble along with a rapid outflux. This process was repeated and the gas bubble continuously grew. As a result, effects of multiple fs laser pulses should not be a simple accumulation of the effect of the single fs pulse.

**Fig. 5.**
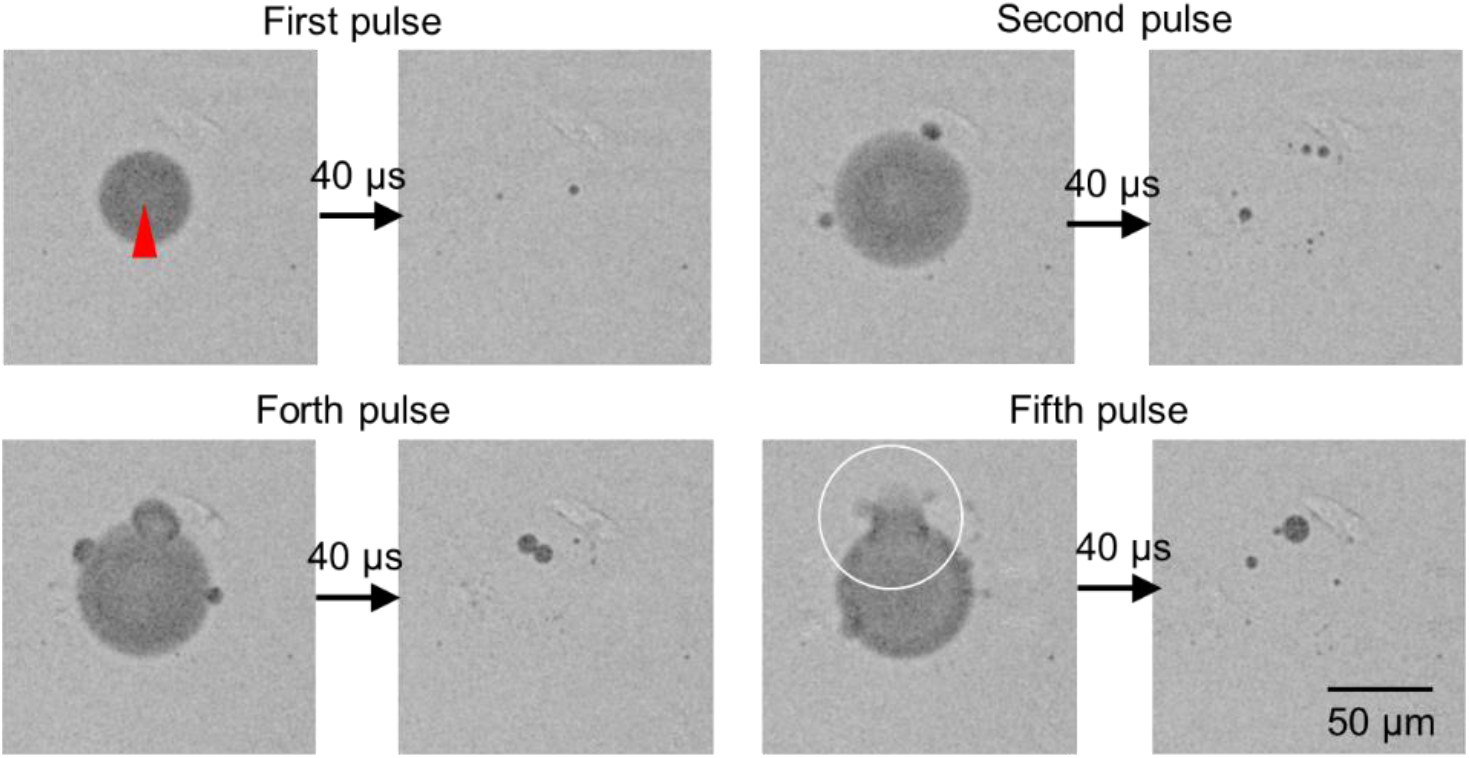
High speed transmission images of cavitation and gas bubble interactions by sequential pulse irradiations at 1 kHz. The images after 1^st^, 2^nd^, 4^th^, 5^th^ pulse were selected. The red triangle indicates the laser focal point. A white circle shows the gas bubble burst after arriving the fifth pulse.

When fs laser pulses were led to a cell sheet at 32 Hz, cell detaching in a wider area than the size of the cavitation bubble were observed, which would be due to strong outflux driven by the cavitation-gas bubbles interaction. Therefore, we have to pay attention to the presence of gas bubbles for laser processing of biological medium with laser pulse irradiations with shorter interval of the gas bubble lifetime.

## 4. Conclusion

Characteristic behaviors of gas bubbles induced by fs laser breakdown were investigated by high speed imaging under different cell culture environments. We concluded that the dissolved air and photochemical decompositions were the main contents of the gas bubbles. The gas bubbles tended to remain near cell probably due to aspherical collapse of the cavitation bubble near boundary. At the same time, the contact of the gas bubbles to a cell membrane did not cause critical damage on its viability. This can be explained that the gas bubble surface was immediately covered and stabilized by amphipathic ingredients in the cell culture medium. However, when multiple fs laser pulses were focused within a lifetime of the gas bubble, interaction with the cavitation bubble led to the gas bubble burst and liquid jet flow, which might cause serious damage on cells.

These results provide important insights for biological tissue processing with intense fs laser pulses. In order to avoid unexpected gas bubble behaviors, fs laser pulse trains with lower frequency and higher pulse energy would be more suitable for the processing rather than higher frequency and lower pulse energy. The limit of this study is that only cell viability is evaluated. The contact of the gas bubble may associate with other physiological effects for example a generation of reactive oxygen species, and a mechanically induced signal transduction. As a future work, these physiological effects must be clarified for better understanding of fs laser processing in biomedical applications.

